# Leading researchers in the leadership of leading research universities: meta-research analysis

**DOI:** 10.1101/2024.04.02.587840

**Authors:** John Ioannidis

## Abstract

It is unknown to what extent leading researchers are currently involved in the leadership of leading research universities as presidents or as executive board members. The academic administrative leader (president or equivalent role) of each of the 146 Carnegie tier 1 USA universities and of any of the top-100 universities per Times Higher Education (THE) 2024 ranking and the members of the executive governing bodies (Board of Trustees, Council, Corporation or similar) for the each of the top-20 universities per THE 2024 ranking were examined for high citation impact in their scientific subfield. Highly-cited was defined as the top-2% of a composite citation indicator (that considers citations, h-index, co-authorship adjusted hm-index and citations to papers as single, first, last authors) in their main scientific subfield based on career-long impact until end-2022 among all scholars focusing in the same subfield and having published ≥5 full papers. Very highly-cited was similarly defined as the top-0.2%. Science was divided into 174 fields per Science-Metrix classification. 38/146 (26%) tier 1 USA university leaders as of end-2023 were highly-cited and 5/146 (3%) were very highly-cited. The respective figures for the top-100 THE 2024 universities globally were 43/100 and 12/100. For the 13 US universities among the top-20 of THE 2024, the probability of their leader being highly-cited was lower (6/13, 46%) than the probability of a randomly chosen active full tenured professor from their faculty being highly-cited (52-77%). Across 444 board members of 14 top-10 THE 2024 universities with data, only 65 (15%) were academics, and 19 (4%) were highly-cited; academics were rare in USA university boards. Board members had predominantly careers in for-profit companies. In conclusion, leading research universities have a dearth of leaders who are high-impact researchers.

## 1. INTRODUCTION

Universities have multiple purposes and missions, including education, public service, and scholarship with discovery, improvement, and dissemination of knowledge. Research is a key component of their missions. The focus and prominence on the research mission may vary across different universities, but it is unquestionably a quintessential consideration in the most prestigious universities worldwide. An important question is whether the leadership of universities, in particular those that have large, intensive, and influential research portfolios, includes leading researchers. Previous work conducted two decades ago suggested that more prestigious universities were more likely to have as leaders people with strong research credentials and citation influence [1–4]. However, evidence has also suggested that very few university leaders were among the most highly-cited scientists worldwide and few of the most highly-cited scientists ventured into university leadership [5].

Given that many years have elapsed since these evaluations and universities have gone through major transformations and challenges in the 21^st^ century [6–8], it would be important to revisit the current situation with recent data and also with better citation tools compared with those available in the past. Currently available data would allow better placement of the citation impact of university leaders across the scientific workforce and also compared with professors in their own institutions. Moreover, previous work had focused on the single leaders in the university academic ladder (presidents or chancellors). While these university leaders are indeed central in university administration, the highest level of executive authority in most universities typically belongs to other governing bodies above them. These executive governing bodies have oversight and fiduciary responsibility for all university affairs and the president or chancellor reports to them. Sometimes the president/chancellor may also be a member of these larger groups. These executive governing bodies are called with different names (e.g. Boards of Trustees, Councils, Corporations) and they are appointed with different mechanisms (e.g., usually by selection of new members by existing members in private universities, by appointment by state authorities in many state universities, or by country leadership in communist countries). There has been no systematic evaluation of whether these executive governing bodies include any members with high citation impact in the scientific literature.

The present evaluation used comprehensive composite indicators to examine the presence of highly-cited and very highly-cited researchers among the leaders of the most prestigious universities in the USA [9] and worldwide [10] and also examined the same features for the members of the governing bodies of the top-20 universities worldwide according to the Times Higher Education (THE) 2024 World University Rankings [10].

## 2. METHODS

### 2.1 Academic university leaders

Evaluations of high ranking according to composite citation indicators, as described below, were performed for the single top academic leader of each of the 146 Carnegie tier 1 USA universities [9] and of any of the top-100 universities per THE 2024 World University Rankings [10]. By default, this was chosen to be the president of the university. However, for public university systems with many campuses, the chancellor of the specific campus was selected (e.g. for the University of California, Berkeley, the chancellor of Berkeley was chosen, instead of the president of the entire University of California system that includes many campuses each with its own chancellor). Chancellors were chosen also for universities where the highest academic leader was called a chancellor and for universities where the president was primarily a ceremonial head (e.g. UK universities). When the position was in transition, the interim holder of the leadership position (interim president, interim chancellor) was chosen. The website of each university was searched to identify the relevant academic leader as of December 20, 2023.

### 2.2 Executive board members

Members of the executive governing bodies (Board of Trustees, Council, Corporation or similar) for the each of the top-20 universities per THE 2024 ranking were also retrieved between December 20, 2023 and February 26, 2024 based on the respective website of each university. It was noted whether the academic leader of the university (president or chancellor, as described above) was included in the membership. For all other members, it was noted how many they were; and how many had a primary academic (or primary research-oriented) background based on the biographies that were available on the university website. For those with primary academic or research-oriented background, evaluations of whether they were highly ranked according to composite citation indicators, as described below, were performed. For the other members, their main listed affiliations/occupations were noted and those mapping to one of the top-100 companies in the Forbes 2023 list [11] were specifically tabulated. Citation or other analyses were not performed when the university website did not contain at least short biographies or bullets listing the main occupation/affiliation that would allow the accurate identification of each member with high confidence.

### 2.3 Databases of composite citation indicators

To calculate composite citation indicators, the Scopus database [12] was used with a data freeze on October 2023, including data as of the end of calendar year 2022 [13]. The database of top-cited scientists is available in public [13] and updated every year. For methods on the development and validation of the composite citation indicators and of the respective database, see previous work [14–16]. In brief, authors who have published at least 5 full papers (those included in the Scopus categories “article”, “review”, and “conference paper”) are considered for ranking. There are approximately 10 million scientists with at least 5 such published papers as of end-2022. Each author is assigned to his/her main scientific subfield, classified according to the Science-Metrix classification [17] that divides science into 174 subfields. For each author, the main field is the one where he/she has published more items during his/her career (until end of 2022). In case of ties, the subfield with the highest ratio of author publications relative to the total number of publications is assigned.

The composite citation indicator considers 6 citation metrics in its calculation: total citations, h-index, co-authorship-adjusted hm index, citations to papers as single author, citations to papers as first or single author, and citations to papers as single/fist/last author during the entire career. Details on the calculation of the composite indicator can be found in [14–16]. The scholars in the top-2% of each subfield are selected. Rankings based on the composite indicators are publicly available both with and without self-citations and the latter were used in the current analysis. However, including self-citations made no material difference. The ranking process adjusts for subject matter, because authors are ranked based on the composite citation indicator across all science and, separately, also specifically within the authors assigned to the same subfield. Those who are ranked among the top-2% of their subfield and those who are among the top-100,000 across all science are included in the database. The composite indicator aims to adjust for co-authorship and author positions which offer a surrogate of relative author contributions, in the absence of detailed, systematic data on contributions for each published paper. The 6 indicators are not totally independent (e.g. total citations correlates with h-index), but cumulatively they capture impact from diverse angles and types of contributions, especially adjusting for multi-authorship and authorship positions in multi-authored articles. They may favor older scholars, since citations accrue over time. If anything, this means that university leaders are likely to be favored, if anything, by these indicators, as they tend to be quite senior when they become leaders.

### 2.4 Definitions of highly-cited and very highly-cited

Highly-cited was defined as being included in the top-2% database for career-long citation impact, as described above. Very highly-cited was defined as being among the top- 0.2% of scholars in the same scientific subfield based on the composite citation indicator for career-long impact.

### 2.5 Comparison with all tenured professors in the same universities

For the 13 USA universities that are among the top-20 of the THE 2024 ranking, information was retrieved from their websites on the number of full, tenured professors that actively serve in them based on their latest available statistics as of January 8, 2024. If this information was not available, the number of tenured appointments (which may include a few associate professors) was captured. If none of this information was clear from the university websites, information was obtained from a centralized database [18]. Some schools (in particular medicine and related health fields such as public health and nursing) in most top universities have mostly non-tenured appointments even for full professor ranks. Non-tenured professors therefore were not counted.

It is very difficult, if not impossible, to directly assess how many of these professors are highly-cited, as defined above, as comprehensive centralized lists of all full tenured professors by name are usually not available in university websites. Therefore, the proportion of active full tenured professors who are highly-cited was approximated indirectly. A random sample of 1000 authors was obtained in Excel from the database of all highly-cited authors. Those with an affiliation with one of the 13 USA universities that are among the top- 20 THE 2024 ranks were identified. For each of them, it was checked online whether they are indeed affiliated or not with that university, and if so, if they are deceased, emeriti, or currently active. Among those currently alive, it was also examined (by perusing information on their names online) whether they are currently full tenured professors or not (e.g. non-tenured professors, professors at lower ranks, adjunct or visiting faculty, non-tenured staff scientists or other). This allowed to estimate the proportion P(h) (and 95% confidence interval thereof) of the 1000 sampled highly-cited authors who are actively tenured full professors at one of these 13 universities. The total number, N(h), of highly-cited active tenured full professors in these 13 universities was then estimated by multiplying the proportion P(h) by the total number of highly-cited scientists in the database (N=204,488).

The proportion of active full tenured professors who are highly-cited across these 13 universities was estimated by dividing N(h) by the sum of active tenured full professors obtained above. This was then compared with the proportion of academic leaders in the same 13 institutions who are highly-cited in their subfields.

### 2.6 Statistical analyses

Analyses are descriptive. Exploratory analyses of differences between groups used Fisher’s exact test. Where appropriate, 95% confidence intervals for proportions were also calculated.

## 3. RESULTS

### 3.1 Highly-cited academic university leaders

Overall, 38 (26%) of the 146 tier 1 USA university leaders as of end-2023 were highly-cited and 5 (3%) of the 146 were very highly-cited. The respective figures for the top- 100 THE 2024 universities globally were 43/100 (43%) and 12/100 (9%). Table 1 lists the highly-cited and very highly-cited academic leaders according to country. As shown, the highest percentages of highly-cited leaders are seen in the United Kingdom (UK). In an exploratory comparison, leaders in UK universities tended to be more commonly highly-cited (exact p=0.0515) and very highly-cited (exact p=0.0255) compared with leaders in other countries. In the USA, leaders of tier 1 universities who were also among the top-100 of THE 2024 tended to be more likely to be highly-cited (16/38, 42% versus 22/108, 20%, exact p=0.0168). Very highly-cited academic leaders focused on very diverse scientific subfields. Among the USA universities’ very highly-cited leaders these subfields were Networks & Telecommunications, Obstetrics & Reproductive Medicine, Polymers, Sport Sciences, and Nanoscience & Nanotechnology. For non-USA universities’ very highly-cited leaders these subfields were Neurology & Neurosurgery (n=2), Psychiatry, Food Science, Networks & Communications, Environmental Sciences, Physical Chemistry, Developmental Biology, and Economics (n=1 each). Given that there are 174 science subfields in the Science Metrix classification, data are sparse to address whether specific subfields are more likely to provide academic leaders, or even more so, highly-cited academic leaders. Moreover, data are too sparse to assess whether there are differences between countries in subfield-specificity for their leaders.

**Table 1.** Highly-cited and very highly-cited academic leaders in top universities according to country.

| Country | Top-universities | Highly-cited academic leader | Percentage | Very highly-cited academic leader |
| --- | --- | --- | --- | --- |
| <u>CARNEGIE</u> |  |  |  |  |
| <u>TIER 1</u> |  |  |  |  |
| USA | 146 | 38 | 26 | 5 |
| <u>THE 2024</u> |  |  |  |  |
| <u>TOP-100</u> |  |  |  |  |
| USA | 38 | 16 | 42 | 3 |
| China and Hong Kong | 12 | 6 | 50 | 2 |
| United Kingdom | 11 | 8 | 73 | 4 |
| Germany | 8 | 3 | 37 | 1 |
| Netherlands | 6 | 3 | 50 | 1 |
| Australia | 5 | 1 | 20 | 0 |
| Canada | 3 | 2 | 67 | 0 |
| Switzerland | 3 | 2 | 67 | 1 |
| France | 3 | 0 | 0 | 0 |
| South Korea | 3 | 0 | 0 | 0 |
| Other* | 8 | 2 | 25 | 0 |

THE: Times Higher Education \*Other includes Japan (n=2), Singapore (n=2), Sweden (n=2), Belgium (n=1) and Russia (n=1)

### 3.2 Comparison of academic leaders against other professors in the same universities

Table 2 lists the number of active tenured full professors in each of the 13 US universities who are among the top-20 of the THE 2024 list and the number of highly-cited scientists with affiliation with any one of these 13 universities in a random sample of 1,000 highly-cited scientists. As shown, in total, these 13 universities account for 7.2 (95% confidence interval, 5.6% to 8.8%) of the highly-cited scientists worldwide. Overall, 28 sampled highly-cited scientists in these 13 universities are clearly active tenured full professors, while the remaining are deceased (n=10), emeritus (n=15), scientists with other ranks or titles (n=6), and medical school full professors for whom it is uncertain if they are also tenured (n=13). Extrapolating to all 204,488 highly-cited scientists, one estimates highly-cited scientists in these universities who are clearly active tenured full professors to be 5,726 (95% confidence interval, 3,681-7,771) and the number would increase up to 8384 (95% confidence interval, 5,930-10,838) if the n=13 sampled full professors at medical schools are also tenured. The sum of active tenured full professors in these 13 universities is 10,928 (Table 2). Therefore, one can estimate that at least 52% (95% confidence interval, 34- 71%) of the active tenured full professors in these universities are highly-cited, and the proportion would increase to 77% (95% confidence interval 54-99%), if the n=13 sampled medical school professors are also tenured. Conversely, only 6 of the 13 (46%) academic leaders of these universities are highly-cited; 2 of these 6 were actually interim presidents at the time of the survey (December 20, 2023).

**Table 2.** Faculty and highly-cited faculty in 13 premier universities in the USA.

| University | Faculty* | Types of faculty counted (source(s) of information considered) | Highly-cited (eligible) Among N=1,000 sample** |
| --- | --- | --- | --- |
| Berkeley | 879 | Full professor rank, all types (https://ofew.berkeley.edu/sites/default/files/faculty_profile_over_time_fall_2023.pdf) | 8 (7) |
| Caltech | 266 | Tenured faculty, full-time (https://www.collegefactual.com/colleges/california-institute-of-technology/academic-life/faculty-composition/) | 2 (1) |
| Columbia | 1010 | Full professor rank, tenured (https://opir.columbia.edu/sites/default/files/content/Statistical%20Abstract/opir_faculty_rank.pdf) | 4 (1) |
| Cornell | 822 | Tenured professorial, full professor rank (https://irp.dpb.cornell.edu/university-factbook/employees) | 4 (2) |
| Harvard | 1068 | Tenured (ladder) (https://faculty.harvard.edu/sites/hwpi.harvard.edu/files/faculty-diversity/files/ay2023_annual_report_-_data_tables.pdf?m=1671128842) | 12 (3) |
| Johns Hopkins | 560 | Tenured faculty full-time (https://provost.jhu.edu/wp-content/uploads/2020/07/2020-Report-on-Faculty-Composition.pdf), 1193 including medicine, nursing and public health (but unclear if all these additional ones would be tenured full-time) | 3 (0) |
| MIT | 704 | Full professor rank (https://facts.mit.edu/employees/faculty/) | 4 (3) |
| Princeton | 541 | Tenured full professor rank (https://dof.princeton.edu/about/facts-figures) | 2 (0) |
| Stanford | 1260 | Tenured, full professor rank (https://facultydevelopment.stanford.edu/data-reports-0/faculty-demographics, https://facts.stanford.edu/academics/faculty-profile/) | 8 (4) |
| U Chicago | 762 | Tenured faculty, full-time (https://data.uchicago.edu/data-employees/, https://www.collegefactual.com/colleges/university-of-chicago/academic-life/faculty-composition/) | 7 (1) |
| U Penn | 863 | Tenured faculty, full-time (https://www.upenn.edu/about/facts, https://www.collegefactual.com/colleges/university-of-pennsylvania/academic-life/faculty-composition/) | 9 (2) |
| UCLA | 1241 | Tenured faculty, full-time (https://apb.ucla.edu/campus-statistics/faculty, https://www.collegefactual.com/colleges/university-of-california-los-angeles/academic-life/faculty-composition/) | 6 (2) |
| Yale | 952 | Full professor rank (ladder, tenured) (https://oir.yale.edu/data-browser/faculty-staff/faculty/faculty-headcounts/faculty-headcount-school-and-rank-w055) , includes medicine, nursing and public health (681 if they were to be excluded) | 3 (2) |
| ALL 13 | 10928 |  | 72 (28) |

\*In counting the number of faculty, the aim to capture the number that reflected active, tenured full professors (or the number closest to this group); medicine, nursing, public health faculty were excluded, unless only tenured professors could be counted \*\*the total number reflects all scientists with an affiliation reflecting the respective university among a random sample of 1,000 highly-cited scientists drawn from the database of 204,488 highly-cited scientists. All affiliations related to the specific university were considered so as to capture also scientists that were listed with an affiliation reflecting a school, center, medical center, or institute related to the university (e.g. Wharton for U Penn or Kennedy School for Harvard). Eligible are those who would also fit the definition used for types of faculty counted in the respective university (generally reflecting those who are active, tenured full professors). Not counted among the eligible are deceased (Columbia 1, Harvard 2, Johns Hopkins 2, Stanford 1, U Chicago 2, U Penn 1, UCLA 1), emeritus (Berkeley 1, Cornell 1, Harvard 1, Princeton 2, Stanford 2, U Chicago 1, U Penn 5, UCLA 1, Yale 1), other ranks/appointments (Caltech 1, Harvard 1, MIT 1, Stanford 1, U Chicago 1, UCLA 1), and medical school full professors for which it is unknown if they are tenured (Columbia 2, Cornell 1, Harvard 5, Johns Hopkins 1, U Chicago 2, U Penn 1, UCLA 1).

### 3.3 Executive board members

As shown in Table 3 information on the occupation of executive board members could be retrieved for 14 of the top-20 THE 2024 universities. Excluding presidents (or similar rank), across 444 board members, only 65 (15%) were academics, and of those only 19 (4%) were highly-cited. Nine of these 14 universities were in the USA and they consistently had very few or no academics among their executive members: across a total of 339 board members, there were only 21 academics (6%) and of those only 7 (2%) were highly-cited scientists. A similar pattern was seen in the National University of Singapore (only 3/20 trustees being academics, none highly-cited). Conversely, in the 3 UK universities, academics comprised a slight majority (40/78, 51%) of council members and 9 (12%) were highly-cited. The executive board of ETH Zurich also had 4 of 7 members being academics and 3 of them were highly-cited.

**Table 3.** Executive board members of top-20 THE 2024 universities: academic members and highly-cited members.

| University | Executive Board | Members excluding president or similar | Academic members (highly-cited) |
| --- | --- | --- | --- |
| Oxford | Council | 27 | 19 (6) |
| Stanford | Board of Trustees | 30 | 0 (0) |
| MIT | The MIT Corporation | 74 | 5 (2) |
| Harvard | Harvard Corporation | 12 | 3 (1) |
| Cambridge | Council | 25 | 13 (3) |
| Princeton | Board of Trustees | 36 | 5 (1) |
| Caltech | Board of Trustees | 78 | 6 (3) |
| Imperial College | Council | 26 | 8 (6) |
| Berkeley | Board of Trustees | 136 | no data |
| Yale | Current Board | 16 | 2 (0) |
| ETH Zurich | Executive Board | 7 | 4 (3) |
| Tsinghua U | Current Administrators | 12 | no data |
| U Chicago | Board of Trustees | 48 | 0 (0) |
| Peking U | Current administration | 14 | no data |
| Johns Hopkins | Board of Trustees | 37 | no data |
| U Penn | Trustees | 48 | no data |
| Columbia | Trustees | 20 | 0 (0) |
| UCLA | Board of Regents | 25 | 0 (0) |
| NUS | Board of Trustees | 20 | 3 (0) |
| Cornell | Board of Trustees | 63 | no data |

Typical occupations of non-academic members of executive boards were leaders of for-profit companies, such as investment funds, other financial sector groups, strategic consulting, law firms, marketplace, real estate, media, entertainment, sports, technology, biomedicine, communication, and others. There was also some lesser representation of leaders of not-for-profit organizations and philanthropists, and occasional members with a track record of government service (including also military).

Table 4 shows for each of the 14 top-20 THE 2024 universities with available data, the companies listed as main occupations/affiliations of board members, limited to the companies that among the top-100 companies in the Forbes 2023 list. There are a total of 27 occurrences pertaining to 18 different companies (detailed information in Supplement 1). 16 of the 39 primarily US top-100 Forbes companies are represented, as opposed to only 2 of 61 top-100 Forbes companies primarily from other countries (p<0.001).

**Table 4.**
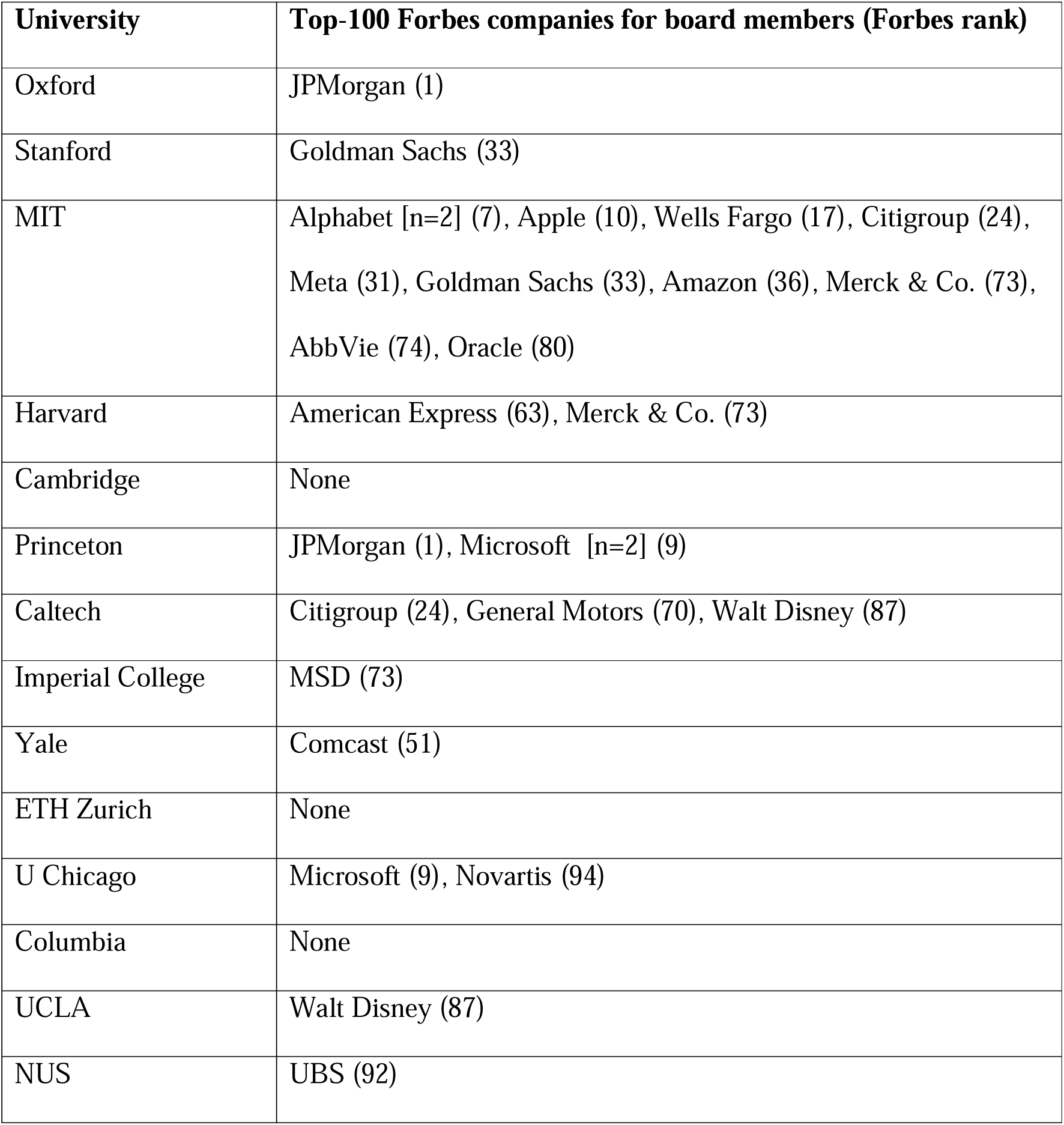
Top-100 Forbes companies listed in main occupations of board members of top universities.

## 4. DISCUSSION

This empirical meta-research analysis shows that even in the leading universities in the USA and worldwide, it is the exception rather than the rule to have university presidents or equivalent academic leaders who are highly-cited in their scientific field. UK universities may have the highest rates of highly-cited scientists serving as academic leaders (typically vice chancellors). Very highly-cited academic leaders are a rarity overall, even more so in American universities. Among the top 13 US universities, the chances that their academic leader is highly-cited are probably lower than the chances of a full tenured professor being highly-cited. Apparently, university presidents on average cannot match the citation credentials of their average faculty, although there is large heterogeneity between different presidents. Furthermore, besides the university presidents, few other academics serve on the executive boards of top universities, especially in the USA, while such participation is more common in UK and Switzerland. Highly-cited academics are almost non-existent in the executive board leadership of top American universities. The executive boards of these universities are comprised largely of businessmen and businesswomen with strong careers in leadership roles among for-profit corporations, including many of those among the Forbes top-100. Of note, top-cited researchers may include some who are currently active in scientific investigation and others who had mostly past accomplishments in research that account for their cumulative citation impact. The overall dearth of top-cited scientists in executive board positions means that these boards lack both scientists who are currently active in what is hot in research and scientists who had a strong past trajectory in scientific investigation.

Most past research on university presidents has used interviews, surveys and document analysis, as shown in a review of 111 studies [19]. A few citation analyses were conducted many years ago [2,5]. The results should be compared with caution given the different citation indicators used. However, it is likely that the under-representation of highly-cited researchers in the university leadership has become substantially worse over time, especially in the USA. For example, 15 years ago, 6 of the 96 academic leaders of 96 high research activity universities were included in the ISI highly cited researchers database [5] that included fewer than 7000 scientists. Conversely, currently only 2 of the academic leaders of these institutions would be among the top-7,000 scientists according to the composite citation indicator database. It is also unknown whether currently an association may exist between the overall university performance and the research track record of its president. Past research has shown that department chairs with higher citation impact also affect the impact of their department after serving some time as chairs [20]. University presidents are more remotely removed from direct research production or even supervision. However, they may be setting a tone for the institution at large. Moreover, they may be seen with skepticism or even cynicism by their faculty, if their scholarship usually places them below the average.

Leading clever people is a difficult enterprise even under the best circumstances [21]. While university communication officers and media allies may try to paint a picture of university presidents as being top scholars, the distance between media manipulation and obvious reality makes these reputation dressing efforts even more repulsive to truly serious scholars.

Admittedly, administration and research scholarship are different roles and may require different skills and perhaps even different personalities. One may thus argue that sound and efficient administrators may still be taken seriously by their academic community, even if they are not highly-cited scholars themselves. Successful single stories can be evoked in defense of this perspective. However, it may an oxymoron to have routinely low- scholarship or no-scholarship leaders lead top scientists. Universities have undergone major transformations in the last 2-3 decades with a major rise of commercialization and managerialism and a disconnect between teaching and research [6,8]. Over time, there have been dramatic changes in the typical profile requested of academic leaders [22]. There may be even more uncertainty of what a university leader should be nowadays in a rapidly changing environment of new challenges and threats for higher education. Moreover, given the time demands of these lucrative, high-salaried positions, even highly-cited scholars are unlikely to be able to maintain their scholarship while serving as presidents. This would lead to their gradual decline from citation rankings and, more essentially, their increasing disconnect with what current research means and what new problems a researcher faces on a day-to-day basis. The disconnect may be even greater when failing to understand the struggles of young researchers and those who are disadvantaged because of structural inequities.

Other researchers who have investigated the profile of university leaders have suggested [1] that it is good to have great scholars running academic institutions.

Nevertheless, one may point out that it does not mean that they have to be the very best. For example, Goodall [1] suggests that presidents should be approximately among the top 10- 20% of scholars. This may not be far from the top-2% group as defined in the current analysis, because the top-2% is captured among the pool all scholars who have published at least 5 full papers. This is a very wide pool that includes not only full professors, but also scholars at all stages of their career development, excluding only the early steps when someone has only published 1-4 papers. Therefore the top-2% in the current analysis may correspond roughly with the top 10-20% of scholars who have reached the career stage of professor or equivalent – and may thus be considered for leadership roles.

While faculty used to run most of the important functions of universities in the past, faculty seem to have almost completely lost control of university executive functions in modern universities [23]. Governing executive boards are the most powerful players in this regard. These governing boards have very few academics among their members, especially in the USA. In fact, most of these already sparse academics come mostly from other institutions. The situation is apparently different in some countries such as UK and Switzerland, where academic faculty continue to serve also in leadership/board roles. What makes a good trustee and a good board in general has been the topic of a considerable literature, but it is unclear how to answer this question for university boards [24,25]. What is clearly obvious, nevertheless, is the complete dominance of successful people with a non-academic, non-research career in for-profit corporations. Some sectors of the economy seem to be more heavily represented but there is large diversity across universities. Top-100 companies have a strong presence in many universities, with the most prominent presence being at the MIT Corporation. Previous research has described the strong interlock between the boards of universities and the boards of powerful corporations [26]. Nevertheless, it is unclear what, if any, flow of information and of shared interests and conflicts this may signify. Information on university websites about trustees is limited. More transparency about processes and rules to protect from conflicts of interest and to enhance accountability within the university and to the public are warranted, given how powerful universities have become on influencing public matters and priorities.

The findings of the current analysis resonate also with some of the power struggles that have recently taken place in several top US universities between board members and faculty. As many top universities favor the very rich, their wealthy board members are likely to be very forthright about what they want. Experienced corporates might think they know better how to run universities, but universities are not (or at least probably should not be) corporations. Even if universities are seen primarily through an angle of management, it is still not clear why they need outside corporates to run them and control them. E.g., the Big 4 firms in management consulting (Deloitte, KPMG International, PricewaterhouseCoopers, and Ernst & Young) would not allow outsiders to control them to the extent that outsiders control our best universities.

Some limitations of the present analysis should be discussed. First, citation metrics have widely known shortcomings [27] and should be used with caution. The ranking of any single scientist may be too favorable or too unfavorable versus their real impact, which may be elusive to capture in full an objectively. However, the overall citation picture of a large number of academic leaders is likely to provide some reasonable, objective benchmark.

Second, some universities may be actively engaged in gaming citation metrics through various spurious efforts, as in the case of Saudi Arabian and Chinese institutions [28–30] and this could affect eventually all citation-based measurements, including those of their leadership. This is less likely to be a problem for most of the universities analyzed here.

However, use of centralized metrics that account also for gaming should be encouraged [31]. University rankings themselves are also subject to gaming and have major shortcomings [32,33], but it is undeniable that the universities analyzed here are leading institutions, even if their exact ranking can be debated. Most scientists would probably agree that the universities assessed here have a high reputation and usually also have high resources to attract excellent scientists. However, this may change with increasing geopolitical tensions. The response of institutional administrative leaders to emerging unprecedented, at times even chaotic, challenges may shape whether these institutions continue to be seen as highly attractive or not.

Third, the current analysis focused on top institutions only. Based on previous analyses [2,5] and the current observation that THE 2024 university ranking correlated with the chances that a university president would be highly-cited, it is almost certain that lower ranked institutions have even fewer highly-cited leaders, on average. Nevertheless, this statement should be tempered by the presence of very high heterogeneity on how academic leaders and executive board members are selected. The processes vary a lot across countries, private versus public versus state-owned (e.g. communist) settings, and local idiosyncratic choices. For example, board members in private institutions may be selecting new members that reflect their own business allies, conflicts and biases. The extent and nature of inbreeding as well as state and political interventions in selecting university leaders needs better study.

Fourth, the current analysis did not consider other leaders at lower leadership levels, e.g. vice presidents. However, vice president positions are typically even more strongly administrative and supportive in nature and it is likely that successful researchers would be even less likely to be attracted to them – with some exceptions.

In conclusion, top research universities worldwide are rarely run by exceptional researchers and executive boards in the USA are almost devoid of exceptional researchers in their membership. This should raise discussion about what universities and their leadership stand for.

## Funding

None.

### Data sharing

All key data are in the manuscript and in databases that are already open to the public.

## SUPPORTING INFORMATION

### Supplement 1. Detailed information for table 4

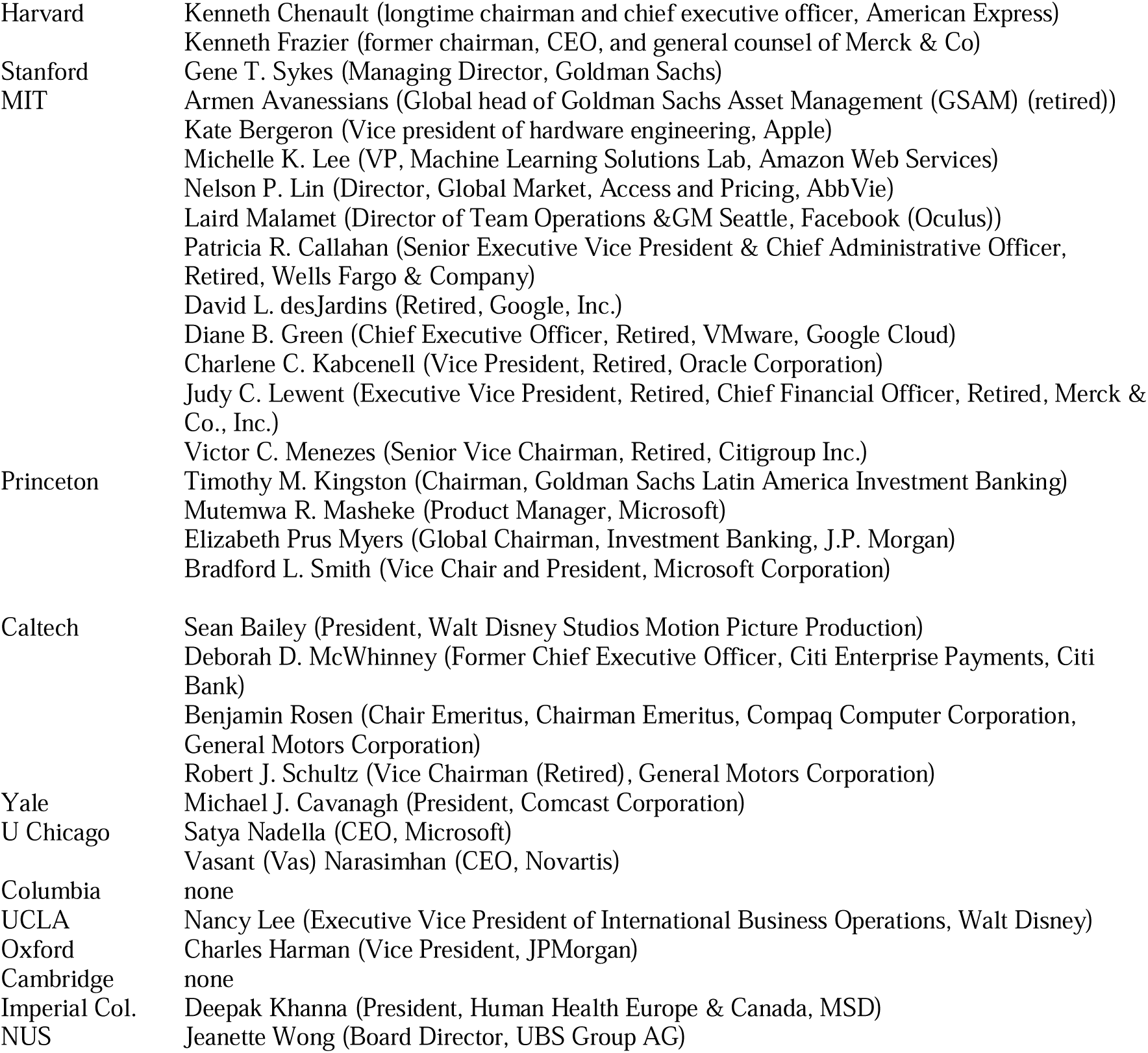

Weblinks for information on trustee members

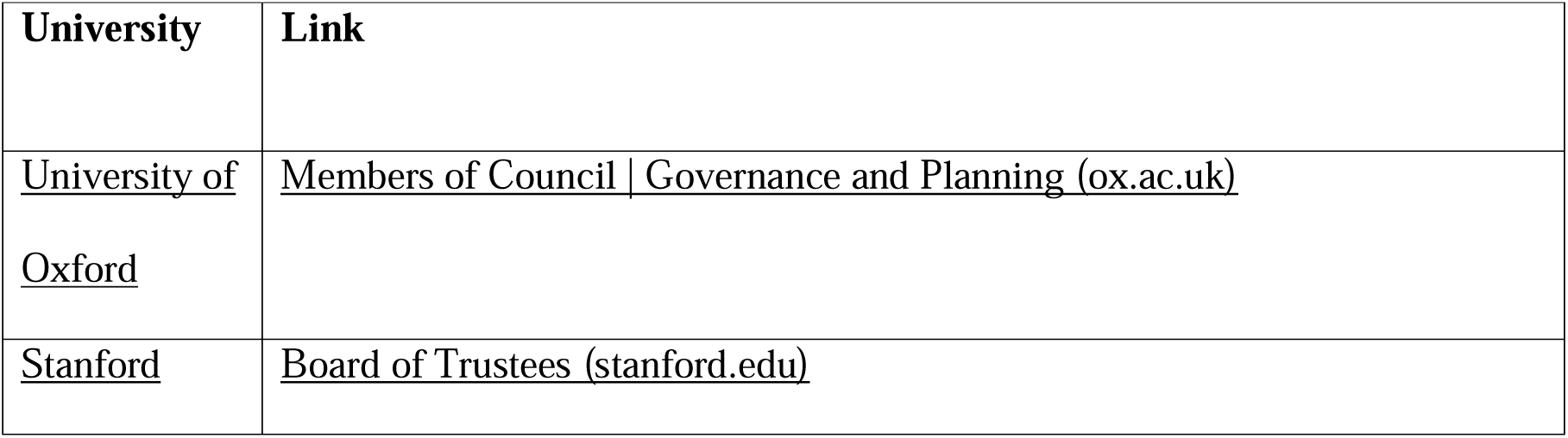
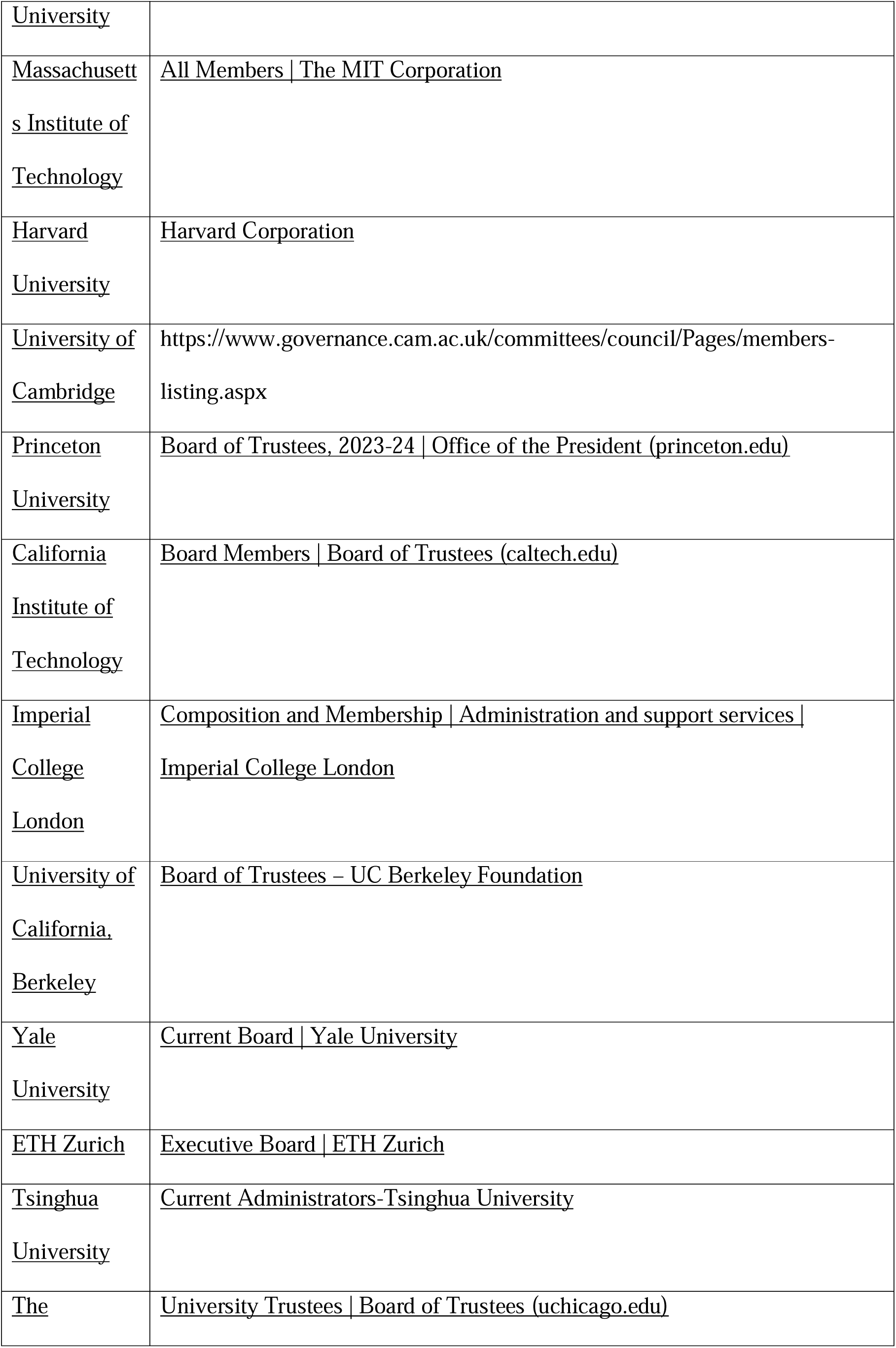
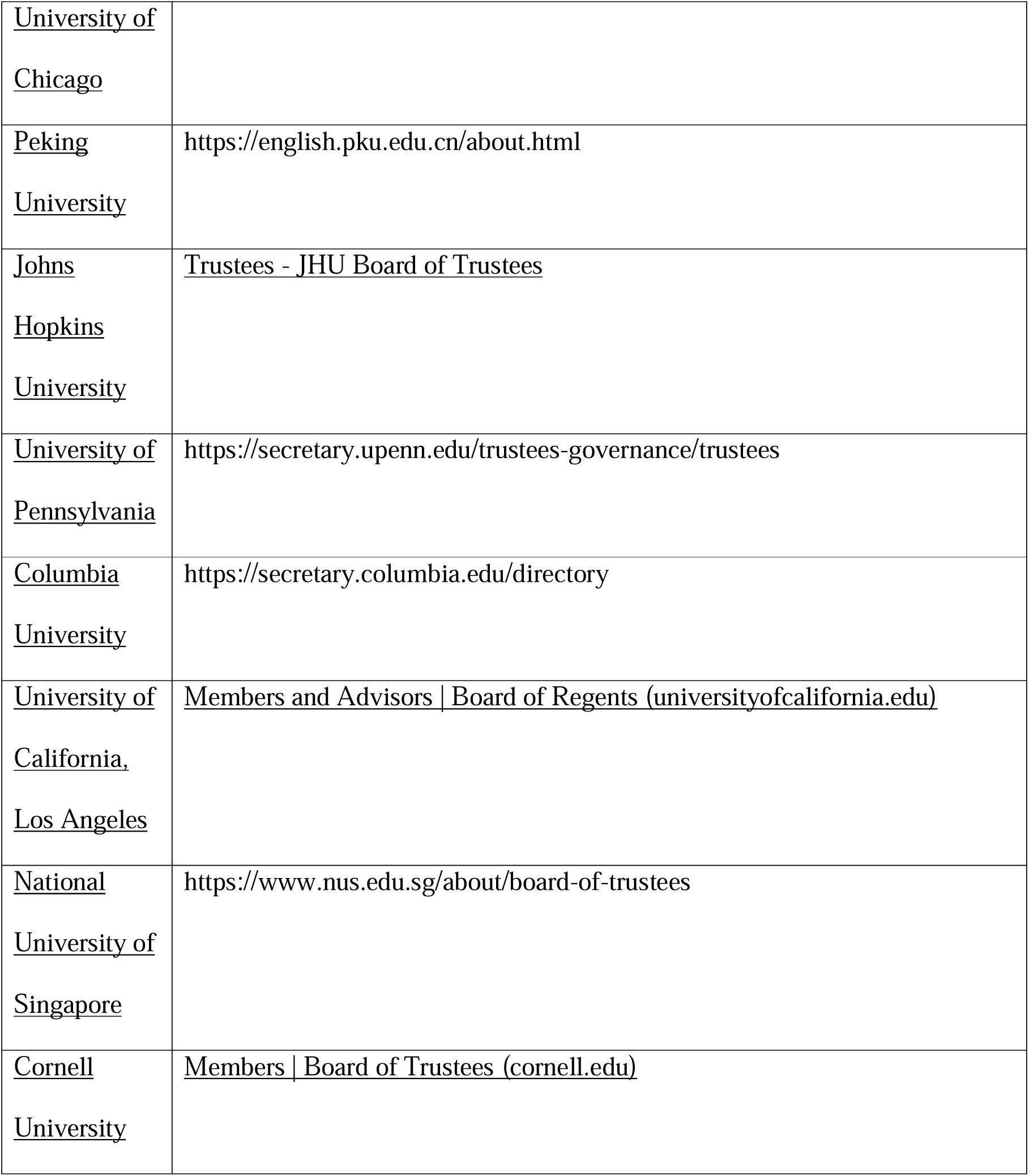

